# Network preservation analysis reveals dysregulated metabolic pathways in human vascular smooth muscle cell phenotypic switching

**DOI:** 10.1101/2022.06.03.494772

**Authors:** R. Noah Perry, Diana Albarracin, Redouane Aherrahrou, Mete Civelek

## Abstract

Vascular smooth muscle cells (VSMCs) are key players involved in atherosclerosis, the underlying cause of coronary artery disease (CAD). They can play either beneficial or detrimental roles in lesion pathogenesis, depending on the nature of their phenotypic changes. An in-depth characterization of their gene regulatory networks can help better understand how their dysfunction may impact disease progression. We conducted a gene expression network preservation analysis in aortic SMCs isolated from 151 multi-ethnic heart transplant donors cultured under quiescent or proliferative conditions. We identified 86 groups of co-expressed genes (modules) across the two conditions and focused on the 18 modules that are least preserved between the phenotypic conditions. Three of these modules were significantly enriched for genes belonging to proliferation, migration, cell adhesion, and cell differentiation pathways, characteristic of phenotypically modulated proliferative VSMCs. The majority of the modules, however, were enriched for metabolic pathways consisting of both nitrogen-related and glycolysis-related processes. Therefore, we explored correlations between nitrogen metabolism-related genes and CAD-associated genes and found significant correlations, suggesting the involvement of the nitrogen metabolism pathway in CAD pathogenesis. for six genes in the nitrogen metabolism pathway. We also created gene regulatory networks enriched for genes in glycolysis and predicted key regulatory genes driving glycolysis dysregulation. Our work suggests that dysregulation of VSMC metabolism participates in phenotypic transitioning, which may contribute to disease progression and suggests that AMT and MPI may play an important role in regulating nitrogen and glycolysis related metabolism in SMCs.

## Introduction

Coronary artery disease (CAD) is the leading cause of death in the United States (US)^1^. While mortality due to CAD has decreased by approximately 50% in the US since the 1980s, the remaining disease burden still has a large socioeconomic impact on our society. Though this decrease may be attributed to therapies that modify CAD risk factors, such as lipid-lowering and anti-hypertensive drugs, these therapies do not target gene expression in the vessel wall where the disease develops. Vascular smooth muscle cells (VSMCs) make up the medial layer of the vessel wall and have been shown to impact every step of atherosclerosis, the underlying cause of CAD^2^.

VSMCs show remarkable plasticity in response to vascular injury. Quiescent VSMCs can shift to a highly proliferative and migratory phenotype that promotes VSMCs to migrate into the intimal layer of the vessel wall. VSMCs in the intimal layer then produce extracellular matrix components and promote fibrous cap stability, therefore, protecting against plaque rupture. As VSMCs undergo phenotypic switching, they have been shown to lose the expression of traditional VSMC marker genes and dedifferentiate into both atheroprotective (fibroblast-like) and atherogenic (macrophage-like) cell types^3^. Macrophage-like VSMCs become pro-inflammatory, releasing cytokines, enhancing the migration of phagocytic cells and accelerating the rate of cell necrosis and plaque growth. Despite intensive research, the mechanisms driving the structural and functional phenotypical transformations of VSMCs are not fully understood. Uncovering how gene expression in VSMCs is being reprogrammed could lead to the discovery of new treatments.

Systems biology approaches have been used to describe the components of the cardiovascular system^4,5^. These approaches postulate that networks of genes, rather than linear pathways, define complex physiological and pathological processes^6,7^. Therefore, we used gene co-expression networks to identify modules that represent highly correlated transcript profiles in both quiescent and proliferative VSMCs to study the properties of the network modules. We then used network-based preservation statistics to quantify within-module topology that are preserved between the two conditions. By focusing on the modules with the lowest preservation, we were able to identify networks of genes, and therefore complex physiological and pathological processes, specific to each phenotypic condition. A deeper investigation of these modules highlighted reprogrammed gene expression profiles that occur during, and potentially drive, VSMC phenotypic transformation. Because loci associated with CAD through genome-wide association studies (GWAS) have been reported to regulate VSMC plasticity^8^, we then mapped genes in CAD GWAS loci to further demonstrate that the dysregulated pathways may be contributing to phenotypic plasticity and disease progression.

## Materials and Methods

### Smooth Muscle Cell Culture and Gene Expression

Smooth muscle cell (SMC) gene expression dataset and donor characteristics have been described in detail elsewhere^9^. Briefly, we cultured aortic SMCs isolated from 6, 12, 64, and 69 of the individuals with East Asian, African, Admixed American, and European ancestries in complete media (containing 5% FBS) until 90% confluence. We then switched to either serum-free media for 24 hours to mimic the quiescent state of SMCs or continued to culture in complete media to mimic the proliferative state of SMCs ^10^. Total RNA was extracted using the RNeasy Micro Kit (Qiagen) and the RNase-free DNase Set. RNA integrity scores for all samples, as measured by the Agilent TapeStation, were greater than 9, indicating high-quality RNA preparations. Sequencing libraries were prepared with the Illumina TruSeq Stranded mRNA Library Prep Kit and were sequenced to ∼100 million read depth with 150 bp paired-end reads at the Psomogen sequencing facility. We trimmed the reads with low average Phred scores (<20) using Trim Galore and mapped the reads to the hg38 version of the human reference genome using the STAR Aligner^11^. We quantified gene expression by calculating the transcripts per million (TPM) for each gene using RNA-SeQC^12^ based on GENCODE v32 transcript annotations. In addition to protein-coding RNAs, we also measured the non-coding RNA since they have been shown to play significant roles in SMC biology^13^. We considered a gene as expressed if it had more than 6 read counts and 0.1 TPM in at least 20% of the samples. RNAseq data is available from GEO with the accession number GSE193817.

### Gene set enrichment analysis

We performed Gene Set Enrichment Analysis (GSEA) on all expressed genes shared between the quiescent and proliferative conditions using GSEA software version 4.1 and predefined gene sets from the Molecular Signatures Database version 7.5^14,15^. A gene set is a group of genes that shares pathways, functions, chromosomal location, or other features. For the present study, we used the Hallmark gene set, C2 curated gene sets including KEGG and Reactome, and C5 ontology gene sets including Gene Ontology (GO) Biological Process and Molecular Function sets. GSEA ranks all of the genes in the dataset based on mean value differences and calculates gene set significance using an enrichment score defined as the maximum distance from the middle of the ranked list. The enrichment score indicates whether the genes contained in a gene set are clustered towards the beginning or the end of the ranked list.

### Differential gene expression and functional enrichment analysis

We included genes with > 6 reads in at least 80% of the samples for both conditions for differential expression analysis using DESeq2^16^. Genes were differentially expressed between proliferative and quiescent conditions when P_adj_ < 1×10^-3^ and log_2_(fold-change)>0.5. To characterize the functional consequences of gene expression changes associated with proliferative and quiescent conditions, we performed Gene Ontology (GO)^17,18^, KEGG^19–21^, Reactome^22^, and Hallmark^23^ gene set enrichment analysis on differentially expressed genes using the anRichment R package^24^.

### Weighted Gene Co-expression Network Analysis

A gene module is a cluster of densely interconnected genes in terms of co-expression. We used Iterative Weighted Gene Co-expression Network Analysis (iterativeWGCNA)^25^, which uses hierarchical clustering and an adjacency matrix, to identify gene modules. The adjacency matrix is defined as the similarity between the i-th gene and j-th gene based on the absolute value of the Pearson correlation coefficient between the profiles of genes i and j. IterativeWGCNA follows the same principles as WGCNA^26^ but re-runs WGCNA iteratively to prune poorly fitting genes resulting in more refined modules compared to WGCNA. Genes that are not assigned to any of the modules are designated to the grey module. Because these genes are not co-expressed, we did not consider them in our analyses.

### Network preservation analysis

We performed preservation analysis on modules constructed using iterativeWGCNA to study their changes across the two cell culture conditions. To determine whether a pathway of genes is perturbed between the proliferative and quiescent conditions, we studied modules whose connectivity patterns are not preserved between conditions as demonstrated by their module preservation statistics. For this analysis we used the summary statistic, medianRank, implemented in the WGCNA R package^26,27^ as a composite module preservation statistic. medianRank is a rank-based measure that relies on observed preservation statistics. medianRank is calculated as the mean of medianRank.density and medianRank.connectivity. Density is the mean adjacency (connection strength) across all nodes in the network. Connectivity is the sum of connection strengths with the other network nodes. To calculate medianRank.density and medianRank.connectivity, for each statistic a in the reference network, we ranked modules in the test network based on the observed values 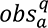. Thus, each module is assigned a rank 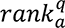 or each observed statistic. The median density and connectivity ranks are then calculated for each module, q, in the test network. The test and reference networks were then flipped to calculate preservation for each condition in the other. A module with a lower medianRank exhibits stronger observed preservation statistics than a module with a higher median rank. We identified the least preserved modules by defining the modules scoring in the bottom 20th percentile of preservation (modules with the highest medianRank score).

### Coronary Artery Disease-associated gene sets

We used two different curated gene sets based on the 175 genomic loci associated with coronary artery disease (CAD) risk in genome-wide association studies (GWAS)^28^. The CAD Candidate gene set includes 2051 genes representing all genes in the 175 CAD GWAS loci. The CAD Prioritized gene set contains 175 genes predicted to be causal at each genome-wide significant loci based on functional annotation, such as genomic location, biological pathway interpretation, literature reviews, and DEPICT gene prioritization^28,29^. Our dataset included 956 genes from the CAD Candidate gene set and 104 genes from the CAD Prioritized gene set.

### Pathway enrichment of co-expression modules

To interpret the biological significance of the least preserved co-expression modules, we performed enrichment analysis based on GO, KEGG, Reactome, and Hallmark pathways using the anRichment R package. We focused on Biological Process and Molecular Function GO terms for downstream analysis.

### Bayesian Network Construction and Key Driver Analysis

We used expression levels of genes identified in co-expression modules as input into the Reconstructing Integrative Molecular Bayesian Networks (RIMBANET) algorithm^30^. One thousand Bayesian networks (BNs) were reconstructed using different starting random seeds. Edges that appeared in greater than 30% of the networks were used to define a consensus network. Edges that were involved in loops were then removed from the consensus network. To identify key regulators for a given regulatory network, we performed key driver analysis (KDA)^31^, which takes as input a set of genes (G) and a directed gene network (N). KDA first generates a sub-network N_G, defined as the set of nodes in N that are no more than h-layers away from the nodes in G. We first computed the size of the h-layer neighborhood (HLN) for each node in the reconstructed BN. For the given network N, *μ* was defined as the average size of the HLN. A score was added for a specific node if the HLN was greater than *μ* + σ(*μ*). Total key driver scores for each node were then defined as the summation of all scores at each h-layer scaled according to h.

### Hypergraph Models

We used PANTHER version 14^32^ to annotate gene ontology terms (GO Term) in BNs. An FDR cutoff of 0.05 was used for enrichment. The HyperG R package was then used to create a hypergraph where each node represents a GO Term and each edge represents a gene or set of genes. The size of the node represents how many genes are in the pathway. Edges and gene labels are color coordinated^33^.

## Results

### Gene expression modules in human smooth muscle cells

We constructed gene co-expression networks from RNA sequencing data of aortic smooth muscle cells (SMCs) isolated from 151 multi-ethnic heart transplant donors cultured in quiescent and proliferative conditions. The overall analysis workflow adopted in this work is summarized in **Figure 1**. After preprocessing and sample outlier detection, 151 samples with gene expression data for 11,330 genes were inputted into the weighted gene co-expression network analysis (WGCNA) ^26^ to create gene co-expression modules for both SMC phenotypic conditions. We performed module detection using iterativeWGCNA^25^. To identify modules of co-expressed genes, we searched for genes with similar patterns of connection strengths to other genes or high topological overlap. A soft threshold power of three and six were used for quiescent and proliferative conditions, respectively, to ensure resulting co-expression networks are closer to a scale-free network frequently observed in large scale biological networks^34^ (**Supplemental Figure 1**). The quiescent condition resulted in 10,764 co-expressed genes segmented into 41 modules, and the proliferative condition resulted in 8,422 co-expressed genes segmented into 45 modules. The modules ranged in size from 34 to 2134 genes. The contingency table in **Supplemental Figure 2** reports the number of genes that fall into quiescent (rows) and proliferative (columns) modules. This table also shows that some modules possess high gene overlap (preserved) across conditions while others appear to be phenotype-specific (unpreserved). Because co-expression modules can capture genes operating within similar biological pathways and functions, deciphering the modules that are context-specific could lead to understanding genes and pathways operating in phenotype-specific context.

**Figure 1.**
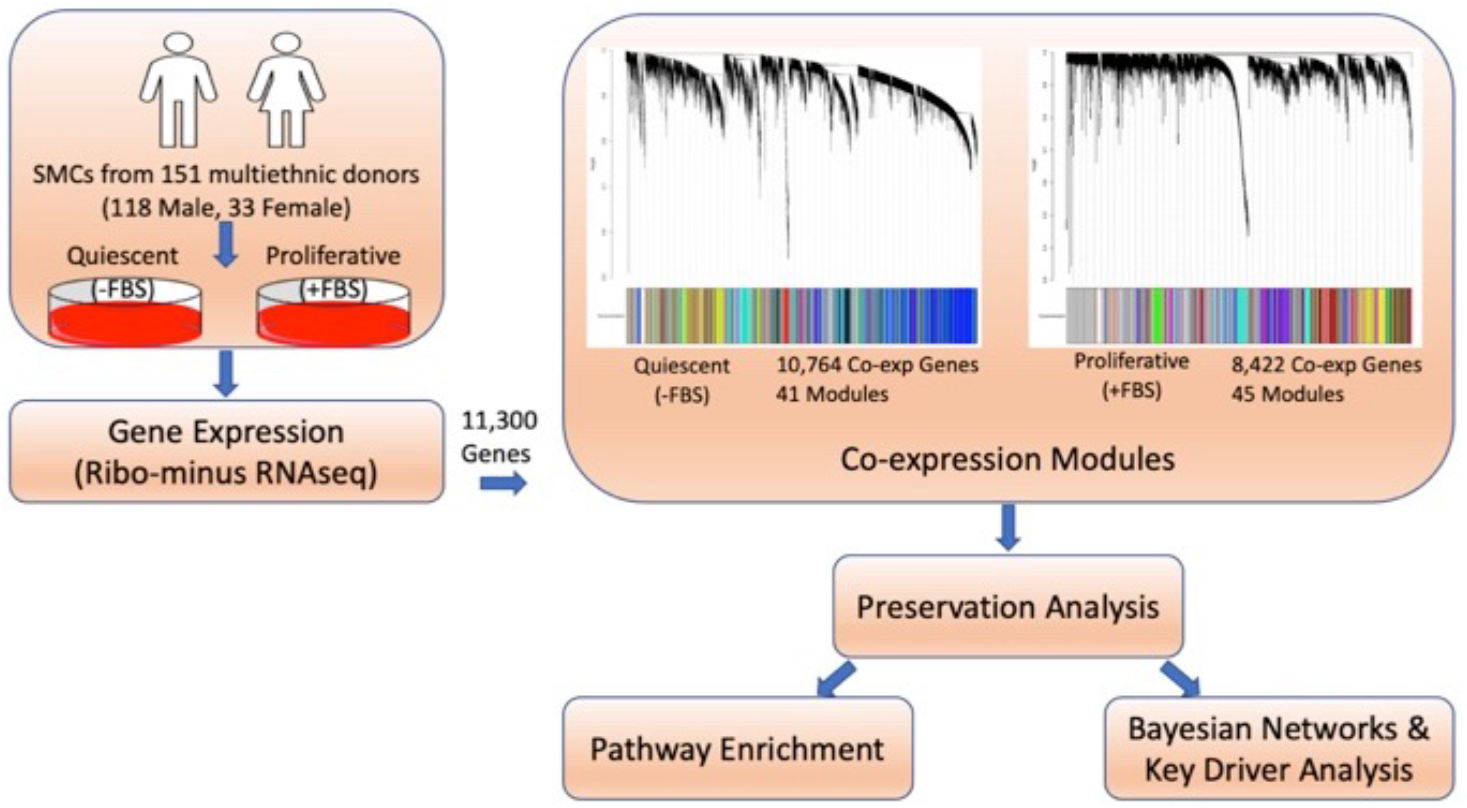
Schematic representation of the overall study design. Smooth muscle cells (SMCs) from the ascending aortas of 151 multiethnic donors were cultured with and without FBS to mimic the quiescent and proliferative phenotypes of SMCs. Gene expression was measured with RNAseq. There were 11,300 genes expressed in at least 80% of the samples across both culture conditions. Co-expression modules were created using the expression levels of the 11,300 genes. Network preservation analysis was performed to rank modules based on preservation. To interrogate the modules, pathway enrichment analysis was performed and Bayesian networks were created. Key driver analysis was performed to identify key regulating genes.

### Preservation analysis of smooth muscle cell modules in distinct phenotypic states

We assessed whether the 41 modules identified in quiescent SMCs were preserved in the 45 modules identified in proliferative SMCs. We utilized statistics that do not depend on a gene’s particular module assignment, but rather network properties such as density and connectivity which rely on connection strengths and topology among all genes^27^. Using a composite statistic of preservation from the WGCNA R package, medianRank, we ranked the preservation of each module across phenotypes. A low medianRank score represented a highly preserved module, whereas a high medianRank represented a less preserved module. To identify the genes and biological pathways most likely to be enriched for phenotype-specific functions, we identified the modules scoring in the bottom 20th percentile of preservation. This cutoff denoted the nine least preserved modules in both the quiescent and proliferative conditions, represented by the modules beneath the red line (**Figure 2A, 2B**). Together these 18 modules contain 2,379 unique genes with topological connectivity representing phenotype-specific interactions. Of the 2,379 genes, less than 10% were shared across the conditions (**Supplemental Figure 3**). Pathway analysis of the 18 modules revealed pathways representative of differential conditions that were not identified with differential gene expression or gene set enrichment analyses (Supplemental Table 1).

**Figure 2.**
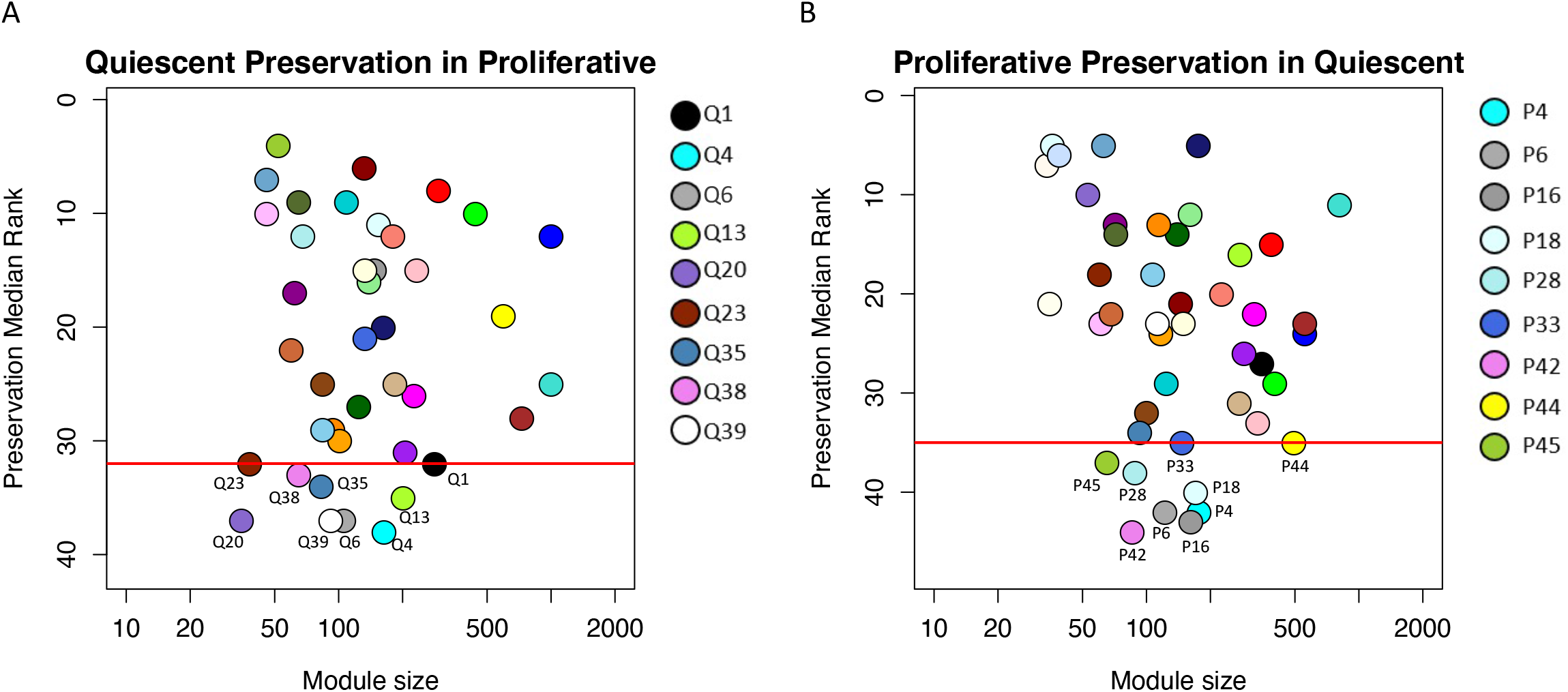
Composite preservation statistics for modules between quiescent and proliferative conditions. The composite statistic, medianRank (y-axis), as a function of the module size. Each point represents a module, labeled by color and a secondary numeric label. Low numbers on the y-axis indicate a high preservation. The red line denotes the bottom 20th percentile of preservation scores. Modules at or below the red line represent the least preserved modules. (A) medianRank scores for the preservation of quiescent modules identified in quiescent SMCs in proliferative modules identified in proliferative SMCs. (B) medianRank scores for the preservation of proliferative modules in quiescent modules. There are nine modules at or below the red line cutoff in each condition.

### Enrichment analysis of unpreserved modules

Multiple unpreserved modules captured previously described biological functions that occur during SMC phenotypic transition from a quiescent to a proliferative state. Gene Ontology (GO) enrichment revealed modules in the proliferative condition to be overrepresented with genes belonging to cell-cell junction organization (P6), vasculature development and migration (P16), and regulation of the SMAD pathway (P42), all representative of the proliferative phenotype^35–37^ (**Figure 3A**). Further, these three modules contain 38 CAD candidate genes and 11 CAD prioritized genes (see Methods). Thirty CAD genes are differentially expressed between quiescent and proliferative conditions, including all 11 prioritized genes (**Figure 3B**). Differential expression of these CAD-associated genes suggests their involvement in the phenotypic plasticity of SMCs.

**Figure 3.**
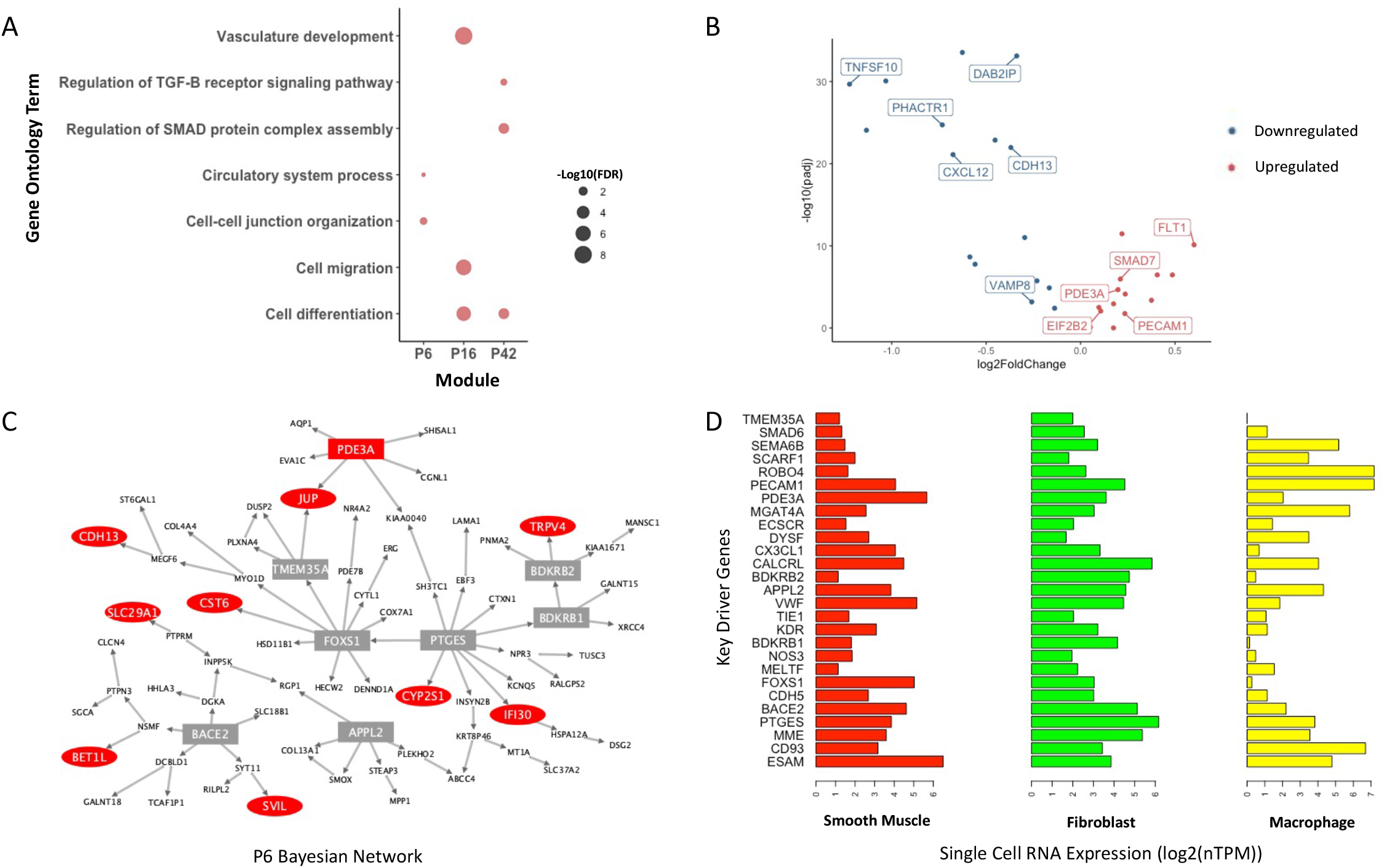
Characterization of modules representative of phenotypic conversion of SMCs from a quiescent to a proliferative state. (A) Gene Ontology enrichment of proliferative SMC related pathways in the P6, P16, and P42 modules. Each point is scaled according to -log_10_(FDR) values. (B) Volcano plot displaying differential expression of the 38 CAD candidate genes present in the P6, P16, and P42 modules. The 11 CAD prioritized genes are labeled. Blue points represent genes downregulated and red points represent genes upregulated in proliferative SMCs compared to quiescent SMCs. (C) Bayesian network created from genes in the P6 module. Red nodes represent CAD candidate genes and square nodes represent genes with a key driver score greater or equal to 1. (D) Horizontal bar plots of single cell RNAseq expression data from Protein Atlas of key driver genes in smooth muscle cells, fibroblasts, and macrophages.

We created Bayesian Networks (BNs) from the co-expression modules to refine regulatory interactions to predict how CAD genes are being regulated and/or regulating gene expression (**Figure 3C, Supplemental Figures 4A, 4B**). We next performed key driver analysis (KDA) to identify the genes with high regulatory potential. We highlighted genes with a key driver (KD) score greater than 1 to capture a wide range of genes with regulatory potential based on network topology. These KD genes are expected to have a greater effect in regulating downstream gene expression and the function of biological pathways. In **Figure 3C**, genes in CAD GWAS loci are denoted by red nodes and genes with a KD score greater than 1 are denoted by a rectangular node. GO Term enrichment analysis of all KD genes across the 3 modules revealed enrichment for cell differentiation (FDR < 0.05), a pivotal function in phenotypic plasticity as VSMCs can dedifferentiate into other cell types, primarily fibroblasts^38^ and macrophages^39^. We then examined RNA single-cell expression profiles of KDs in disease-associated cell types from the Human Protein Atlas^40^. We found that all KD genes are expressed in the dedifferentiated cell types of fibroblasts and/or macrophages providing further evidence that these modules are representative of VSMCs that have transitioned out of a quiescent state into a phenotype representative of atherosclerotic behavior (**Figure 3D**). Together these analyses show strong evidence that using preservation statistics can capture biologically accurate activity occurring in SMCs during the transition from a quiescent to a proliferative state.

### Metabolic pathway enrichment in unpreserved modules

Over half of the unpreserved modules representative of phenotype switching were enriched for metabolic pathways. Of these 10 modules, nine showed enrichment for nitrogen-specific metabolism. Changes in nitrogen metabolism have been demonstrated in endothelial cells (ECs) to promote EC phenotypic transition from a quiescent to a proliferative state^41^. It is well documented that changes in nitrogen content, specifically nitric oxide, also regulate SMC proliferation, migration, and calcification^42,43^, but it is unclear if changes in nitrogen processes are a byproduct or causal of phenotypic switching in VSMCs. 114 of the CAD candidate genes were members of the nitrogen metabolism enriched modules. To further test the potential role of nitrogen metabolism in contributing to phenotypic plasticity, we calculated correlations between expression levels of 8 genes in the Nitrogen Metabolism KEGG pathway expressed in quiescent and proliferative SMCs and 114 genes in the CAD candidate gene set present in our nine nitrogen metabolism enriched modules (**Figure 4B, Supplemental Figure 5**). Seventy-five percent of the Nitrogen Metabolism KEGG pathway genes were strongly correlated (Pearson r >|0.3|, Bonferroni corrected P-value < 5 x10^-5^) with CAD candidate genes in both the quiescent and proliferative conditions. The strongest correlations for both phenotypes were with AMT, which encodes the aminomethyltransferase enzyme. Eleven CAD candidate genes in the quiescent condition and 12 CAD candidate genes in the proliferative condition were strongly correlated with AMT ((Pearson r >|0.5|). AMT has previously been shown to be associated with CAD risk^44^. AMT was also differentially expressed between quiescent and proliferative conditions, suggesting that gene expression levels of CAD GWAS genes functioning in nitrogen metabolic enriched modules were reprogrammed due to changes in AMT expression levels, or vice versa. These data suggest that nitrogen metabolism plays a role in the progression of CAD, potentially through regulating SMC plasticity.

**Figure 4.**
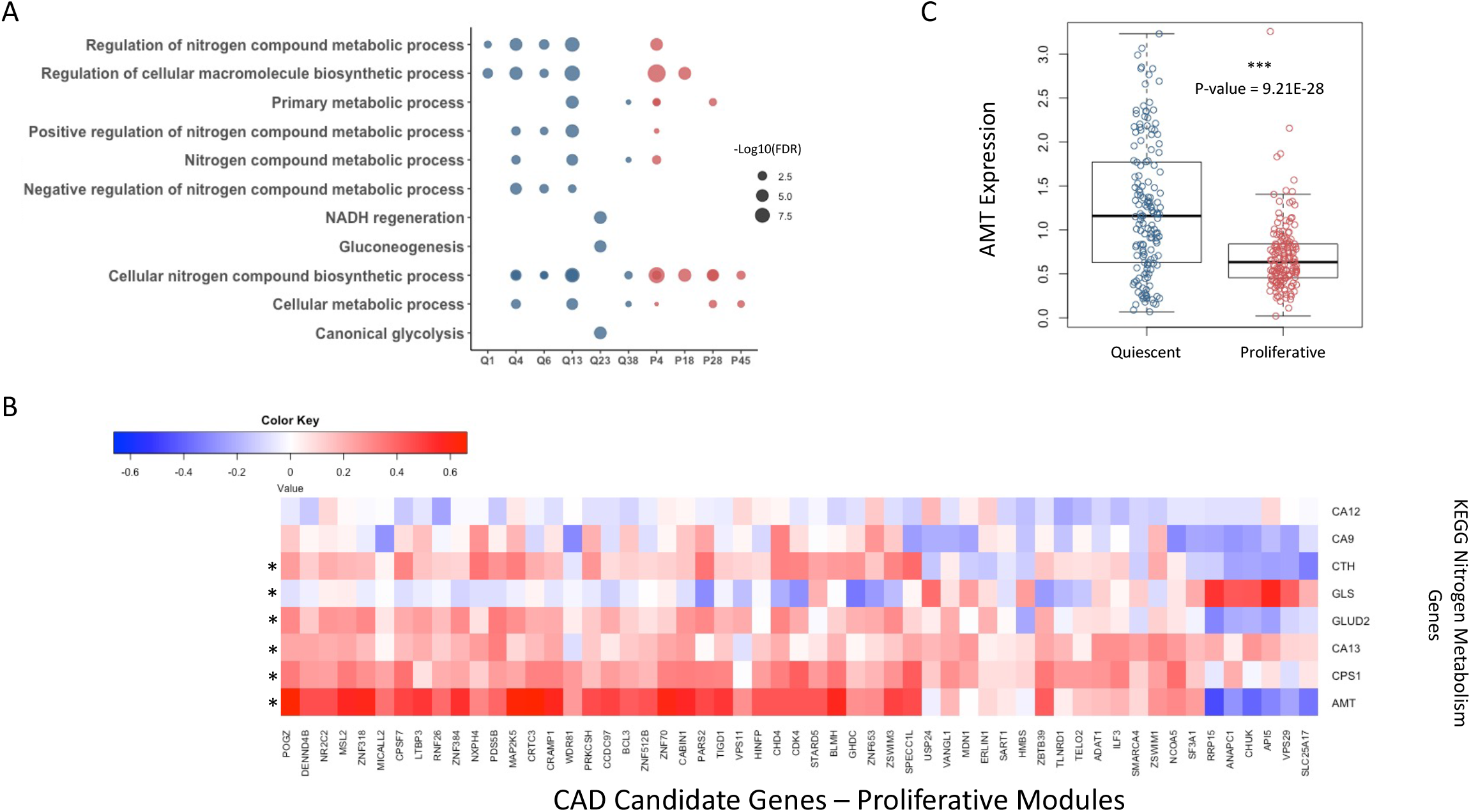
Presence of nitrogen metabolic processes in the least preserved modules. (A) Gene Ontology enrichment of metabolically related pathways in 10 of the least preserved modules. Each point is scaled according to -log_10_(FDR) values. Blue points represent modules from the quiescent condition and red points represent modules from the proliferative condition. (B) Heatmap of Pearson correlations (r) between 55 CAD candidate genes in the P4, P18, P28, and P45 modules and 8 genes in the KEGG Nitrogen Metabolism pathway. Asterisk marks denote genes in the KEGG Nitrogen Metabolism pathway with at least one correlation (r) ≥ |0.3| at a Bonferroni corrected p-value ≤ 5 x10^-5^. (C) Expression levels of the AMT gene in quiescent and proliferative conditions. AMT is downregulated in proliferative conditions (P < 9.3 x10^-28^).

In addition to modules enriched for nitrogen metabolic processes, another module was enriched for NADH regeneration and canonical glycolysis pathways (Q23) (**Figure 4A**). Glycolysis plays an important role in the proliferation of VSMCs^45,46^. Consequently, proliferative VSMCs demonstrate an increase in glycolytic flux so it is unsurprising that we identified a context-specific function of glycolysis^47^. Exploring the glycolytic alterations of VSMCs, however, may provide new insights into the genes involved in the quiescent and proliferative functions of glycolysis. To address the potential rewiring of glycolysis, we compared network topology between the unpreserved Module Q23 in the quiescent condition, and a proliferative module, Module P17, that is also enriched for NADH regeneration and canonical glycolysis. We created BNs for each co-expression module to predict genetic regulatory function and identified KDs to isolate potential genes responsible for driving glycolytic rewiring (**Figures 5A, 5B**). There were 11 shared genes between the two BNs, 7 unique to the quiescent condition, and 11 unique to the proliferative condition. The BNs shared two CAD candidate genes, ENO2 and SPAG4, with the addition of RAB20 in the proliferative network. RAB20 expression was downregulated in proliferative SMCs (P-value <0.001). Downregulation of RAB20 has been shown to promote glycolysis and contribute to enhanced cell proliferation and motility^48^. KD analysis identified a novel KD gene in the proliferative BN, MPI. GO Term analysis of the genes present in each BN showed that all enriched pathways present in the quiescent condition were preserved in the proliferative condition. However, in the proliferative condition, there were more genes in each shared pathway and the addition of new pathways (**Figures 5C, 5D**). Three of the new GO Term pathways in the proliferative BN were represented by the presence of the KD, MPI, suggesting that mannose metabolism could be driving glycolysis rewiring during SMC transition.

**Figure 5.**
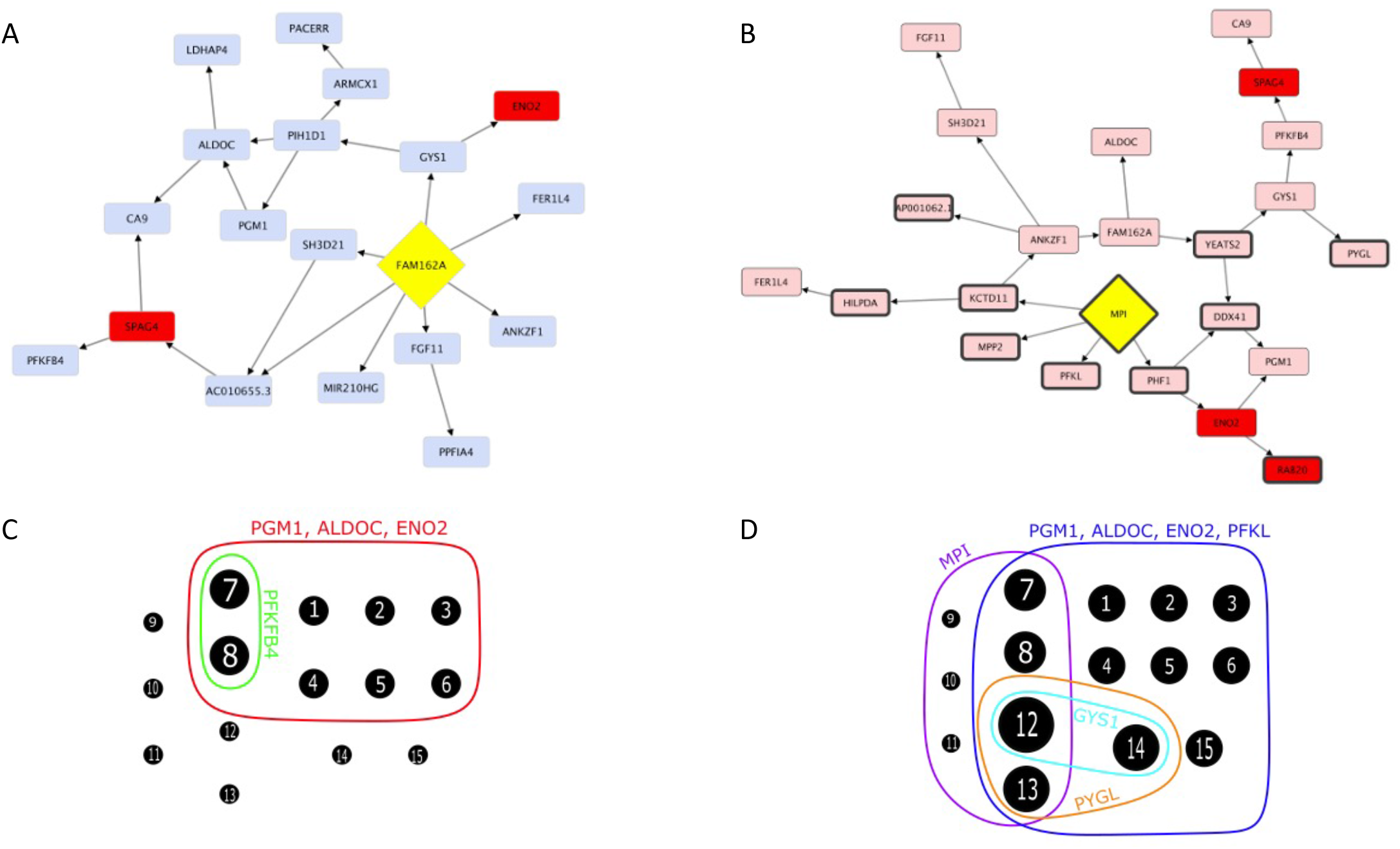
Rewiring of glycolysis metabolic pathway in SMC phenotypic transition. Bayesian networks of genes in the (A) Q23 module and (B) P17 module. Red nodes represent CAD candidate genes and yellow diamonds represent the highest scoring key driver gene. Bold node outlines in (B) represent genes unique to the P17 BN. (C-D) Hypergraph representations of enriched GO Terms (FDR < 0.05) based on genes present in the (C) Q23 and (D) P17 BNs. Each node represents a GO Term. Nodes are scaled according to the number of genes functioning in the GO Term. Edges represent the gene or genes functioning in the surrounding nodes (GO Terms). GO Terms for each node: 1. Glycolytic process 2. ATP generation from ADP 3. ADP metabolic process 4. Purine ribonucleoside diphosphate metabolic process 5. Nucleoside diphosphate phosphorylation 6. Pyruvate metabolic process 7. Hexose metabolic process 8. Monosaccharide metabolic process 9. Mannose-6-phosphate isomerase activity 10. GDP-mannose biosynthetic process 11. GDP-mannose metabolic process 12. Nucleobase containing small molecule metabolic process 13. Carbohydrate metabolic process 14. Ribose phosphate metabolic process 15. Generation of precursor metabolites and energy.

## Discussion

SMCs are a major cell type present at all stages of an atherosclerotic plaque. Previously thought to only play a protective role in stabilizing the fibrous cap, lineage-tracing studies have highlighted that SMCs can adopt alternative phenotypes that positively and negatively contribute to disease progression^49–51^. Investigating the genetic architecture of quiescent SMCs compared to proliferative SMCs could identify biological pathways being rewired during phenotypic transformation, leading to mechanistic predictions driving SMC plasticity in atherosclerosis, the underlying cause of CAD.

Since it is impossible to study these processes in detail in the arteries of living humans, we cultured VSMCs isolated from an ethnically-diverse population of 151 heart transplant donors in two conditions that are believed to represent the quiescent and proliferative state of the cells.

Differential gene expression analysis confirmed the gene expression profiles that are consistent with the phenotypic state of the cells^9^. To our knowledge, this work is the first to propose a comprehensive approach exploring gene co-expression networks observed in proliferative SMCs that are not preserved in quiescent SMCs, and vice versa. Because co-expression networks are representative of functionally related genes, a network preservation approach was able to capture dysregulated pathways whose gene-gene interactions were rewired as a result of phenotypic transition of SMCs

Preservation analysis identified the least preserved gene co-expression modules between quiescent and proliferative SMCs. Three of these modules were significantly enriched for biological pathways representative of SMCs in atherosclerotic lesions, such as proliferation, migration, cell differentiation^3^, cell-cell junction assembly^35^, and SMAD regulation^37^. Capturing physiologically relevant *in vivo* biology assured that our *in vitro* experimental design was able to capture aspects of VSMC plasticity. Further, over half of the unpreserved modules were enriched for metabolic function. Emerging evidence has shown that the metabolism of VSMCs is correlated with the phenotype switching and the progression of atherosclerosis, among other vascular diseases^52^. Unpreserved modules enriched for metabolic functions were present in our quiescent and proliferative conditions representing genetic rewiring of metabolic pathways contributing to a phenotype-specific role of metabolism. Thus, our results support the claims that metabolism of VSMCs are correlated with phenotype switching.

Although nitrogen and glycolytic metabolism in VSMCs are not fully understood, previous reports have identified their potential role in VSMCs and atherosclerosis^52^. We are the first to hypothesize mannose metabolism as a possible mechanism contributing to proliferative VSMCs. Mannose is not a significant energy source in humans but it is required for protein glycosylation^53^. Mannose treatment was shown to attenuate weight gain, improve glucose and lipid homeostasis, and reduce gene expression of inflammatory markers in adipocytes of high-fat diet mice^54^. Mannose supplementation in chow-fed mice, however, did not have as pronounced of an effect on weight gain attenuation suggesting that mannose has a greater impact in pathological conditions. In addition, plasma levels of mannose have recently been shown to be a biomarker of CAD and a more vulnerable plaque phenotype^55^. It is not clear whether mannose is related to CAD because it marks insulin resistance or because of an intrinsic biological property^56^. As VSMCs respond to vascular injury and transition to a more proliferative, disease-like phenotype, mannose metabolism may be responding to a pathological-like phenomenon and mediating changes in metabolism due to imbalances in energy uptake, thus contributing to disease development through a discrete biological mechanism.

This study demonstrates the power of network preservation statistics to identify differences between two biological states. We provide new evidence supporting the role of nitrogen metabolism as a potential regulator of VSMC plasticity. Further studies need to be conducted to discern whether dysregulated metabolism in VSMCs is a byproduct or a driving mechanism of phenotypic plasticity. Specifically, considering the role of AMT in regulating nitrogen metabolism and MPI in regulating mannose and glycolysis metabolism in VSMCs.

