## Supplementary Material for "Network preservation analysis reveals dysregulated metabolic pathways in human vascular smooth muscle cell phenotypic switching"

### Supplemental Tables and Figures

**Supplemental Table 1:** Comparison of the top 25 enriched pathways for methods of Gene Set Enrichment analysis, pathway analysis of differentially expressed genes, and pathway analysis of the 18 least preserved modules. The ranking includes enriched pathways from the Hallmark, Reactome, KEGG, and Gene Ontology databases.

| Database | Gene Set Enrichment Analysis (GSEA) | FDR | Database | Differential Expression | FDR | Database | Least Preserved Modules | FDR |
|---|---|---|---|---|---|---|---|---|
| REACTOME | G2/M Checkpoints | 0.01 | HALLMARK | E2F Targets | 1.00E-18 | GO TERM | RNA Processing | 3.82E-20 |
| KEGG | Pyrimidine Metabolism | 0.01 | REACTOME | Cell Cycle | 4.99E-17 | GO TERM | Vasculature Development | 7.17E-14 |
| REACTOME | Cell Cycle Checkpoints | 0.011 | REACTOME | Cell Cycle, Mitotic | 1.36E-16 | GO TERM | Cardiovascular System Development | 8.86E-14 |
| REACTOME | RHO GTPases Activate Formins | 0.011 | HALLMARK | G2M Checkpoint | 2.32E-14 | GO TERM | Blood vessel Development | 1.03E-13 |
| REACTOME | Extension of Telomeres | 0.012 | GO TERM | Mitotic Cell Cycle | 1.77E-13 | HALLMARK | Hypoxia | 1.64E-13 |
| REACTOME | Cell Cycle Mitotic | 0.013 | GO TERM | Cell Cycle Progress | 2.44E-13 | GO TERM | Transcription Regulator Activity | 3.64E-12 |
| HALLMARK | MYC Targets V1 | 0.013 | GO TERM | Mitotic Cell Cycle Process | 3.39E-13 | GO TERM | Blood Vessel Morphogensis | 5.71E-12 |
| REACTOME | Metabolism of Nucleotides | 0.013 | GO TERM | Cell Cycle | 8.59E-12 | GO TERM | Anatomical Structure Formation Involved in Morphogenesis | 2.08E-11 |
| REACTOME | Mitotic G1 Phase and G1/S Transition | 0.013 | GO TERM | DNA-dependent DNA Replication | 8.59E-12 | GO TERM | Tube Morphogenesis | 3.29E-11 |
| REACTOME | RHO GTPase Effectors | 0.015 | REACTOME | Activation of the Pre-replicative Complex | 9.34E-12 | GO TERM | Angiogensis | 3.56E-11 |
| REACTOME | Homology Directed Repair | 0.017 | REACTOME | Activation of ATR in Response to Replication Stress | 4.51E-11 | GO TERM | Tube Development | 7.33E-10 |
| REACTOME | S Phase | 0.017 | GO TERM | DNA Replication Initiation | 2.76E-10 | GO TERM | Positive Regulation of RNA Metabolic Process | 1.47E-09 |
| REACTOME | Cell Cycle | 0.018 | GO TERM | DNA Replication | 2.99E-10 | GO TERM | Regulation of Transcription by RNA Polymerase II | 2.23E-09 |
| REACTOME | Processing of DNA Double Strand Break Ends | 0.018 | REACTOME | Mitotic Prometaphase | 3.38E-09 | GO TERM | Transcription by RNA Polymerase II | 2.52E-09 |
| REACTOME | M Phase | 0.019 | GO TERM | Cell Division | 1.78E-08 | HALLMARK | Glycolysis | 2.66E-09 |
| HALLMARK | G2M Checkpoint | 0.02 | REACTOME | Resolution of Sister Chromatid Cohesion | 1.76E-08 | GO TERM | Anatomical Structure Morphogenesis | 3.10E-09 |
| HALLMARK | Glycolysis | 0.02 | GO TERM | Nuclear Division | 2.32E-08 | GO TERM | Positive Regulation of RNA Biosynthetic Process | 3.44E-09 |
| REACTOME | DNA Disulfide Strand Break Repair | 0.021 | REACTOME | DNA Strand Elongation | 2.54E-08 | GO TERM | Positive Regulation of Nucleic Acid-templated Transcription | 3.44E-09 |
| REACTOME | G2/M DNA Damage Checkpoint | 0.021 | REACTOME | Unwinding of DNA | 2.54E-08 | GO TERM | Regulation of Cellular Biosynthetic Process | 6.96E-09 |
| KEGG | Purine Metabolism | 0.021 | GO TERM | Chromosome Segregation | 6.86E-08 | GO TERM | Circulatory System Development | 8.57E-09 |
| REACTOME | Telomere Maintenance | 0.021 | GO TERM | Organelle Fission | 7.99E-08 | GO TERM | Cell Communication | 2.00E-08 |
| HALLMARK | E2F Targets | 0.023 | GO TERM | Double-Strand Break Repair via Homologous Recombinations | 1.75E-07 | GO TERM | Locomotion | 2.31E-08 |
| KEGG | Progesterone Mediated Oocyte Maturation | 0.023 | GO TERM | Nuclear DNA Replications | 1.85E-07 | GO TERM | Transcription, DNA-templated | 3.15E-08 |
| KEGG | Cell Cycle | 0.025 | GO TERM | Recombinational Repair | 2.05E-07 | GO TERM | Regulation of RNA Metabolic Process | 3.57E-08 |
| REACTOME | L1CAM Interactions | 0.025 | HALLMARK | Cholesterol Homeostasis | 2.61E-07 | GO TERM | Regulation of Trascription by RNA Polymerase II | 5.15E-08 |

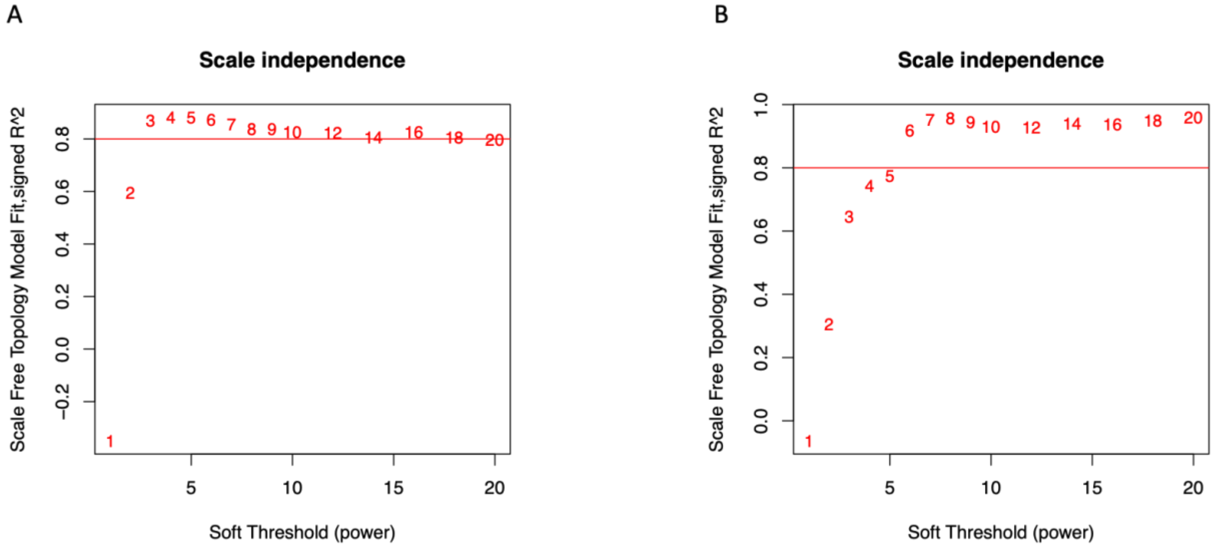


**Supplemental Figure 1:** Determination of soft-thresholding power in Weighted Gene Co-expression Network Analysis. The scale-free fit index (y-axis) as a function of the soft thresholding power (x-axis) for gene expression of (A) quiescent and (B) proliferative smooth muscle cells. An $R^2$ value of 0.8 was used as the cutoff corresponding to scale-free topology.

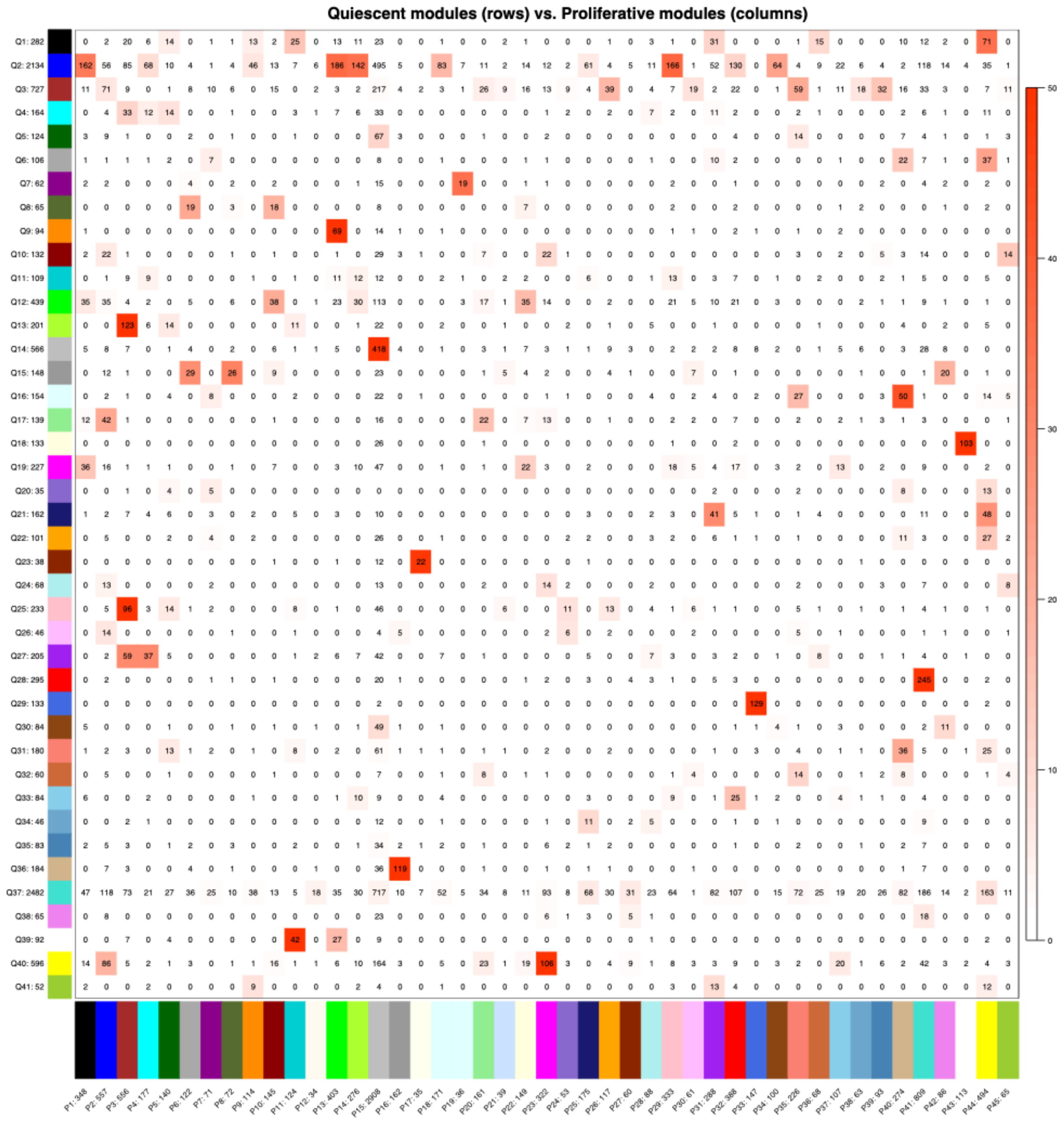


**Supplemental Figure 2:** Overlap table of genes shared in modules across quiescent and proliferative conditions. Cross tabulation of quiescent modules (rows) and proliferative modules (columns). Each row and column is labeled by the corresponding module color and total number of genes in the intersection of the corresponding row and column module. The table is color-coded by the Fisher exact test p-value of the overlap of gene module membership (-log(p)), according to the color legend on the right.

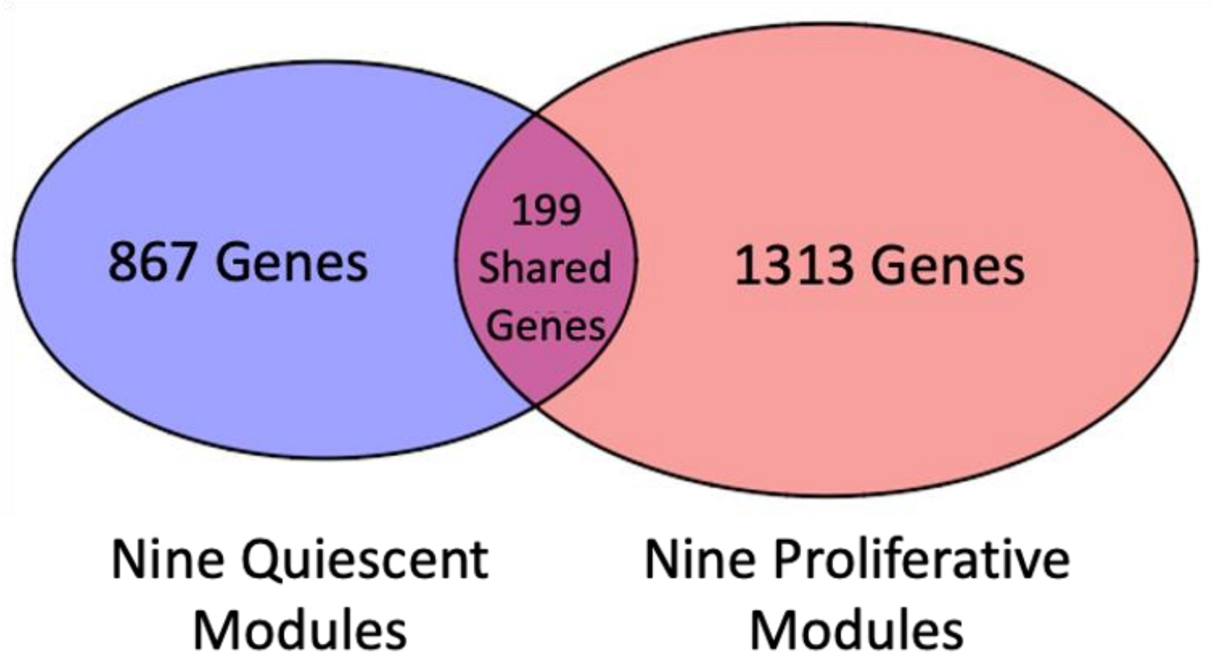


**Supplemental Figure 3:** Representation of genes in the least preserved modules. A Venn diagram of the genes contained in the 18 least preserved modules.

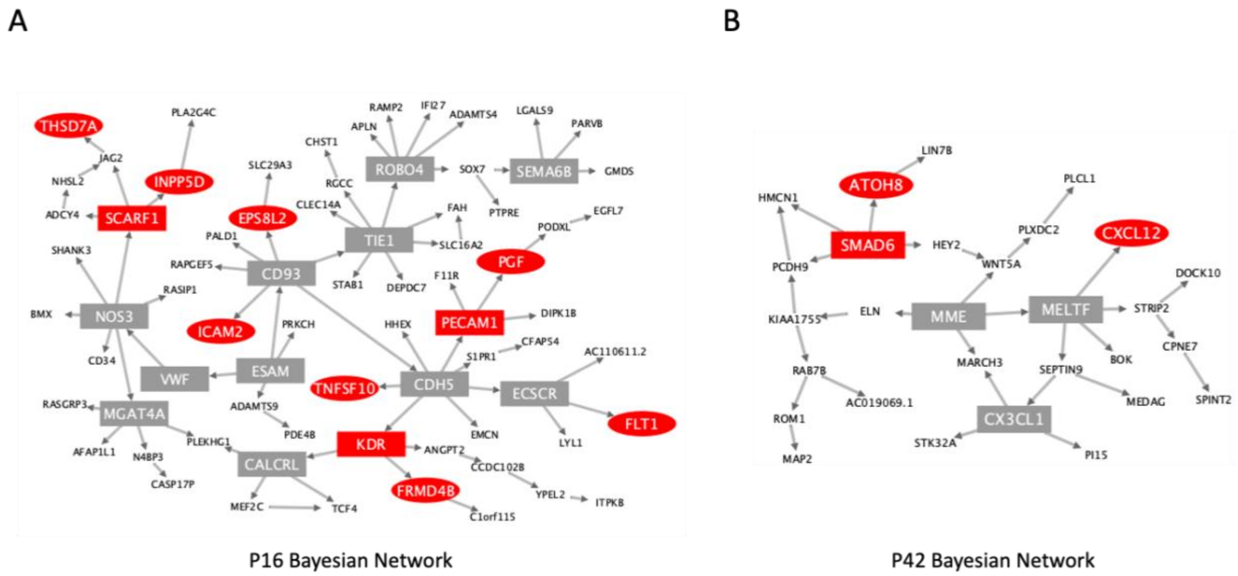


**Supplemental Figure 4:** Bayesian network analysis. Bayesian networks created from genes in the (A) P16 and (B) P42 modules. Red nodes represent CAD candidate genes and square nodes represent genes with a key driver score greater or equal to 1.

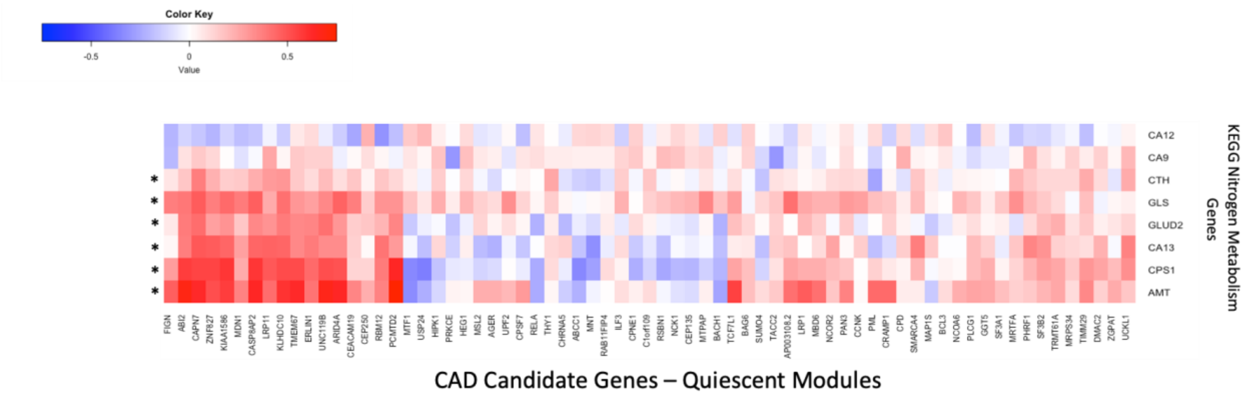


**Supplemental Figure 5:** Correlations between nitrogen related genes and CAD associated genes in quiescent VSMC gene expression. Heatmap of Pearson correlations (r) between 69 CAD candidate genes in the Q1, Q4, Q6, Q13, and Q38 modules and 8 genes in the KEGG Nitrogen Metabolism pathway. Asterisk marks denote genes in the KEGG Nitrogen Metabolism pathway with at least one correlation (r) $\geq |0.3|$ at a Bonferroni corrected p-value $\leq 5 \times 10^{-5}$.
